# Multivariate analysis of multimodal brain structure predicts individual differences in risk and intertemporal preference

**DOI:** 10.1101/2024.07.04.602046

**Authors:** Fredrik Bergström, Guilherme Schu, Sangil Lee, Caryn Lerman, Joseph W. Kable

**Affiliations:** Faculty of Psychology and Educational Sciences, University of Coimbra, Portugal; Department of Psychology, University of Gothenburg, Sweden; Social Science Matrix, University of California, Berkeley, CA, USA; Department of Psychiatry, Perelman School of Medicine, University of Pennsylvania, Philadelphia, PA 19104, USA; Department of Psychology, University of Pennsylvania, Philadelphia, PA 19104, USA

**Keywords:** magnetic resonance imaging (MRI), brain structure, thresholded partial least squares (T-PLS), risky choice, intertemporal choice

## Abstract

Large changes to brain structure (e.g., from damage or disease) can explain alterations in behavior. It is therefore plausible that smaller structural differences in healthy samples can be used to better understand and predict individual differences in behavior. Despite the brain’s multivariate and distributed structure-to-function mapping, most studies have used univariate analyses of individual structural brain measures. Here we used a multivariate approach in a multimodal data set composed of volumetric, surface-based, diffusion-based, and functional resting-state MRI measures to predict reliable individual differences in risk and intertemporal preferences. We show that combining twelve brain structure measures led to better predictions across tasks than using any individual measure, and by examining model coefficients, we visualize the relative contribution of different brain measures from different brain regions. Using a multivariate approach to brain structure-to-function mapping that combines across many brain structure properties, along with reliably measured behavior phenotypes, may increase out-of-sample prediction accuracies and insight into neural underpinnings. Furthermore, this methodological approach may be useful to improve predictions and neural insight across basic, translational, and clinical research fields.

---

Historically, it was only possible to make inferences about brain structure and behavior by studying individuals with brain damage (Yu, Kan, & Kable, 2020). However, the advent of magnetic resonance imaging (MRI) has made it possible to non-invasively investigate relationships between brain structure and behavior in healthy samples. Many brain-wide association studies have linked brain structure to behavioral traits, but with mixed results, likely in part because of small sample sizes (Mdn = 23; Marek et al., 2022) and using univariate analysis of individual brain structure measures (Genon et al., 2022; Makowski et al., 2023; Song et al., 2022). Here we used a multivariate approach in a multimodal MRI data set to predict two reliably-measured behavioral traits – risk and intertemporal preferences.

Risk preferences (i.e., the tendency to choose larger risky versus smaller certain rewards) and intertemporal preferences (i.e., the tendency to choose larger later versus smaller sooner rewards) are two moderately stable individual differences (Chuang & Schechter, 2015; Escobar et al., 2023; Hertwig et al., 2019; Mata et al., 2018; Meier & Sprenger, 2015; Seneca et al., 2012; Zeynep Enkavi et al., 2019) linked to important outcomes in life. Risk preferences have been associated with heavy drinking, smoking, being overweight/obese, seat belt non-use, not having insurance, holding stocks instead of treasury bills (Anderson & Mellor, 2008; Barsky et al., 1997; Lejuez et al., 2003, 2005); self-employment status (Ekelund et al., 2005), pathological gambling (Branas-Garza et al., 2007; Wiehler & Peters, 2015), and financial decisions (Noussair et al., 2014). Intertemporal preference has been associated with heavy drinking (Vuchinich & Simpson, 1998), smoking (Bickel et al., 2008; Yi & Landes, 2012), pathological gambling (Wiehler & Peters, 2015), drug addiction (MacKillop et al., 2011), and excessive credit card borrowing (Meier & Sprenger, 2020). Understanding the relationship between brain structure and risk or intertemporal preferences is therefore an important endeavor.

Several studies have tried to link structural brain measures to risk or intertemporal preferences (for review, see Kable & Levy, 2015). Regarding risk preferences, a greater preference for larger risky over smaller certain rewards has been positively associated with grey matter volume (GMV) in right posterior parietal cortex (Gilaie-Dotan et al., 2014a; Grubb et al., 2016a; Quan et al., 2022); amygdala (Jung et al., 2018; Quan et al., 2022); and cerebellum (Quan et al., 2022); negatively associated with cortical complexity, gyrification, and sulci depth in ventromedial prefrontal cortex (vmPFC) and gyrification in dorsomedial prefrontal cortex (dmPFC) (Bergström et al., 2024); and associated with both structural and functional connectivity between vmPFC and amygdala (Jung et al., 2018). Moreover, Jung et al. (2018) found that a model with amygdala GMV, functional amygdala-mPFC connectivity, and amygdala-mPFC white matter tract strength together explained more variance than any measure alone. Furthermore, white matter coherence between anterior insula and anterior ventral striatum was associated with a reduced preference for positively skewed gambles (Leong et al., 2016).

Regarding intertemporal preferences, a greater preference for smaller immediate over larger delayed rewards has been negatively associated with GMV in dorsolateral prefrontal cortex (dlPFC) and inferior frontal cortex (Bjork et al., 2009); ventral putamen (Cho et al., 2013), superior frontal gyrus (Schwartz et al., 2010); insula, middle temporal gyrus, and entorhinal cortex (Owens et al., 2017); vmPFC (Bergström et al., 2024; Owens et al., 2017); anterior temporal cortex (Garzón et al., 2023); and positively associated with GMV in vmPFC and anterior cingulate cortex (Cho et al., 2013), ventral striatum (Schwartz et al., 2010), and posterior cingulate cortex (Cho et al., 2013; Schwartz et al., 2010); negatively associated with cortical thickness in vmPFC (Bergström et al., 2024; Bernhardt et al., 2014; Drobetz et al., 2014; Pehlivanova et al., 2018); temporal pole and temporoparietal junction (Pehlivanova et al., 2018); and entorhinal cortex (Lempert et al., 2020); negatively associated with cortical complexity and gyrification in vmPFC, and gyrification and cortical thickness in dmPFC (Bergström et al., 2024); negatively associated with white matter tract strength and functional connectivity between dlPFC and striatum, and positively with the same measures between amygdala and striatum (van den Bos et al., 2014); negatively associated with “white matter integrity” (i.e., negatively with fractional anisotropy (FA) and positively with mean diffusivity) in the frontal cortical-striatal tract (Peper et al., 2013) and lateral prefrontal cortex (Olson et al., 2009); positively associated with functional resting-state connectivity between ventral striatum and vmPFC (Costa Dias et al., 2013; Li et al., 2013), between dorsal anterior cingulate cortex and anterior insula (Li et al., 2013), and between vmPFC and fronto-insular cortex (Han et al., 2013); and negatively associated with functional resting-state connectivity between dlPFC and parietal cortex (Li et al., 2013).

However, these previous studies linking brain structure to risk and intertemporal preferences have primarily relied on univariate analyses of single structural brain measures. In contrast to univariate approaches, multivariate approaches can combine multiple brain measures and account for the relationships between brain structural properties and brain areas. Many different brain areas and brain structural properties are likely associated with any specific behavior, so using multimodal imaging and multivariate analyses may therefore increase power and insight (Genon et al., 2022; Song et al., 2022). However, few studies have used multimodal imaging and multivariate analyses to link brain structure and behavior (but see, Han, Gu, Brown, Zhang, & Liu, 2020; Rasero, Sentis, Yeh, & Verstynen, 2021). Here we performed multivariate brain structure-to-function mapping with Thresholded Partial Least Squares (T-PLS; Lee et al., 2022), combining several structural, diffusion, and functional resting-state MRI measures to predict two reliably measured behavioral phenotypes, risk and intertemporal preferences, and visualize the relative contribution of each brain measure and brain area.

## Materials and Methods

### Participants

In this study we used a previously published multimodal neuroimaging data set (Jung et al., 2018), which is a subset of data acquired for the Retraining Neurocognitive Mechanisms of Cancer Risk Behavior (RNMCRB) study (Kable et al., 2017a). RNMCRB participants were randomized to receive 10 weeks of adaptive cognitive training (Lumosity games) or non-adaptive, untargeted cognitive stimulation (simple computerized video games), and underwent pre- and post-intervention brain scans. The study acquired high-resolution T1-weighted anatomical MRI, resting-state (RS-)fMRI, diffusion tensor imaging (DTI), and task-based fMRI during two economic tasks to measure risky and intertemporal preferences. Participants were excluded if they had a history of brain injury, a history of psychiatric or substance disorders, current use of psychotropic medication, current use of chewing tobacco, snuff, or smoking cessation products, left-handedness, intellectual disability (< 90 score on Shipley’s Intelligence Quotient (IQ) Test). All study procedures were approved by the Institutional Review Board of the University of Pennsylvania, and all participants provided written informed consent. The main trial outcomes have been described in a previous report but found no effect of cognitive training relative to active control on brain activity, decision-making, or cognitive performance (Kable et al., 2017b).

Of the full cohort at baseline (n = 166), 145 had T1-weighted MRI, RS-fMRI, and DTI images, and 37 additional participants were excluded for low DTI quality (signal-to-noise ratio, n = 17), excessive head motion (> 2 SD from group mean, n = 15) in rs-fMRI data, outliers in risk preference (> 3 SD from group mean; n = 2) and node strength in their graph analysis (> 3 SD from group mean, n = 3) by Jung et al. (2018). Additionally, we checked T1-weighted image quality in CAT12 by performing overall correlations between all participants (to check sample homogeneity) and found three participants with an overall correlation value less than two standard deviations from the median. At closer inspection, these three participants had motion artifacts in their T1-weighted images and were therefore excluded. We therefore used 105 participants (61 males and 44 females; age, M = 24.30, SD = 4.67 years; IQ, M = 111.34, SD = 6.66; intertemporal discount rate (k), M = -1.81, SD = 0.44; risk tolerance (α), M = -0.19, SD = 0.18) for our analyses.

### Risk preferences

In the risky choice task, participants had to choose between receiving a smaller but certain reward (i.e., 100% probability to receive $20) or a larger but riskier reward (e.g., 47% probability to receive $84) on 120 trials. The certain reward was always fixed at $20 on all trials, while the probability and risky reward amount varied across trials. Each trial began by presenting the probability and reward amount for the risky alternative (the fixed certain option was never displayed). Participants had 4 s to accept or reject the risky alternative, after which a marker indicating their choice (a checkmark if the risky alternative was accepted, and “X” if rejected) appeared for 1 s. We estimated individual risk tolerance (α) by fitting the following logistic function to our choice data using maximum likelihood estimation (goodness-of-fit: Mdn_R_^2^ = 0.37, SD = 0.12 with function minimization routines (fmincon) in MATLAB (for code see, https://github.com/sangillee/UMm):

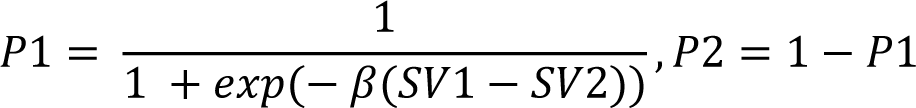

where P1 refers to the probability the risky option was chosen and P2 the probability that the safe option was chosen. SV1 refers to the subjective value of the risky option and SV2 to the subjective value of the safe option. β refers to a scaling factor that was fitted for each subject. The SV of the choice options was assumed to follow a power utility function:

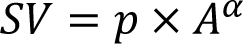

where p is the probability of winning amount A, and α is a risk tolerance parameter that was fitted for each participant. For the risky option, there is always a 1 - p chance of not winning anything. Higher α indicates higher risk tolerance or lower risk aversion.

### Intertemporal preferences

In the intertemporal choice task, participants had to choose between receiving a smaller but immediate reward (i.e., receiving $20 today) or a larger but delayed reward (e.g., receiving $40 in 31 days) on 120 trials. The immediate reward was always fixed at $20 on all trials, while the delay time and delayed reward amount varied across trials. Each trial began by presenting the delay time and reward amount of the delayed option (the fixed immediate option was never displayed). The participants had 4 s to accept or reject the delayed alternative, after which a marker indicating the choice (a checkmark if the delayed alternative was accepted, and “X” if rejected) appeared for 1 s. We estimated individual discount rate (k) by fitting the following logistic function to our choice data using maximum likelihood estimation (goodness-of-fit: Mdn_R_^2^ = 0.58, SD = 0.39) with function minimization routines (fmincon) in MATLAB:

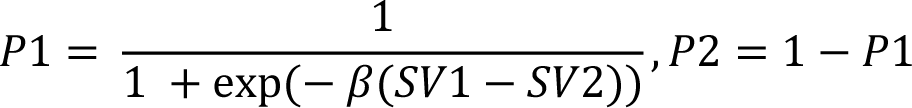

where P1 refers to the probability the delayed option was chosen and P2 the probability that the immediate option was chosen. SV1 refers to the subjective value of the delayed option and SV2 to the subjective value of the immediate option. β refers to a scaling factor that was fitted for each subject. The SV of the choice options was assumed to follow hyperbolic discounting:

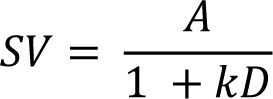

where A is the amount of the delayed option, D is the delay time until receiving the reward (for immediate choice, D = 0), and k is a discount rate parameter that was fitted for each participant. Higher values of k indicate reduced tolerance for delay or greater discounting of delayed rewards.

Both the risky and intertemporal choice tasks were incentive compatible. At the end of the experiment, participants received the option they chose on one randomly selected trial across both choice tasks. If the randomly selected trial was from the risky choice task and the participant chose the risky option, the gamble was resolved by the roll of a die. If the randomly selected trial was from the intertemporal choice task and the participant chose the delayed option, they only received the delayed amount after the number of days specified on that trial. All payments were delivered on a prepaid debit card given to the participant.

### MRI acquisition

A Siemens 3T Trio scanner (Siemens, Erlangen, Germany) was used to acquire MRI data. T1-weighted images were acquired using a magnetization-prepared rapid gradient echo (MPRAGE) sequence [repetition time (TR) = 1630 ms, echo time (TE) = 3.11 ms, voxel size = 0.94 x 0.94 x 1.0 mm^3^, 160 axial slices, 192 x 256 matrix]. Diffusion images were acquired using single-shot spin echo EPI sequences (TR = 8000 ms, TE = 82 ms, voxel size = 1.88 x 1.88 x 2mm^3^, 70 interleaved slices, GRAPPA factor = 3, 30 diffusion directions with b-values of 1000 s/mm2 and 1 image with b = 0 s/mm2). RS-fMRI data were collected using an echo planar imaging (EPI) sequence (TR = 3000 ms, TE = 25 ms; voxel size = 3 x 3 x 3 mm^3^; 53 interleaved axial slices with no gaps; 160 volumes during 8 min and 6 s). During RS-fMRI scanning, participants were asked to keep their eyes open and focus on a fixation cross.

### Preprocessing of MRI data

We used the default processing pipeline of the Computational Anatomy Toolbox (CAT12; (Gaser & Dahnke, 2016) for SPM12 (Welcome Trust Centre for Neuroimaging, London, UK), on Matlab R2019a (Mathworks, Inc., Sherborn, MA, USA), to process the anatomical T1-weighted images and extract average grey matter volume, cortical complexity (i.e., fractal dimension), sulci depth, gyrification index, and cortical thickness for each atlas area.

For voxel-based preprocessing, T1-weighted images underwent spatial adaptive non-local means (SANLM) denoising filter (Manjón et al., 2010), were internally resampled, bias corrected, and affine-registered, followed by standard SPM unified tissue segmentation into grey matter, white matter, and cerebral spinal fluid (Ashburner & Friston, 2005). The resulting brain images were then skull-stripped, parcellated into left and right hemispheres, subcortical areas, and cerebellum; subject to local intensity transformation of all tissue classes to reduce effects of higher grey matter intensities before the final Adaptive Maximum A Posteriori (AMAP) tissue segmentation (Rajapakse et al., 1997), and refined by a partial volume estimation (Tohka et al., 2004). For each participant, grey matter volume was estimated and averaged for each atlas region of the Neuromorphometrics atlas in native space.

For surface-based preprocessing, a projection-based thickness method was used to estimate and reconstruct cortical thickness of the central surface (Dahnke et al., 2013), after which topological correction was applied to repair defects with spherical harmonics (Rachel Aine Yotter et al., 2011), and surface refinement. The final central surface mesh was used to estimate fractal cortical complexity values based on spherical harmonic reconstructions (Yotter et al., 2011), sqrt-transformed sulci depth was extracted based on the Euclidian distance between central surface and its convex hull, and gyrification index was extracted based on absolute mean curvature (Luders et al., 2006). For each participant, cortical complexity, sulcus depth, gyrification index, and cortical thickness values were averaged for each atlas region of the HCP-MMP surface atlas (Glasser et al., 2016) in native space.

For the resting-state fMRI preprocessing, we used the default preprocessing pipeline with Conn Toolbox (Whitfield-Gabrieli & Nieto-Castanon, 2012) to realign and unwarp, slice-time correct, identify outliers, segment & normalize, and denoise. Functional data is realigned with SPM12 realign and unwarp procedure where all volumes are realigned and resampled to the first volume of first run using b-spline interpolation. Field inhomogeneities are estimated and used for susceptibility correction as part of the unwarp step. Temporal misalignment between slices of functional data is corrected with SPM12 slice-time correction by being time-shifted and resampled using sinc-interpolation to match middle of acquisition time. Potential outliers are detected by observing the global BOLD signal and amount of head-motion in scanner. Functional and structural data are normalized into MNI space and segmented into grey matter, white matter, and CFS using SPM12 unified segmentation and normalization procedure. Factors identified as potential confounding effects are estimated and removed separately for each voxel and for each participant and functional run/session using ordinary least squares regression. Potential confounds controlled for were the anatomical component-based noise correction procedure (aCompCor) with noise components from cerebral white matter and CFS (Behzadi et al., 2007), head-motion parameters, outlier scans or scrubbing, constant and first-order linear effects.

Diffusion-weighted images were preprocessed with the FDT diffusion module from the FMRIB (Functional Magnetic Resonance Imaging of the Brain’s diffusion toolbox) software library (FSL, version 6.0.4; Jenkinson et al., 2012) and the MRtrix3 software (Tournier et al., 2019). At each preprocessing step, the images were visually inspected for quality control. Firstly, images were converted from DICOM to NIFTI. For each participant the MPPCA technique (Veraart et al., 2016) implemented in MRtrix3 was used for denoising images. Next, using tools from FSL diffusion toolbox, a whole-brain mask was generated for each subject using their non-weighted diffusion images (b0s), followed by the application of motion and eddy current distortions corrections. Preprocessed images were then diffusion tensor fitted using the least-square technique. To characterize the diffusion displacement in the microstructures of the white matter, the diffusion tensor-based measures fractional anisotropy (FA), mode of anisotropy (MA), and radial diffusivity (RD) were computed. For each participant, each generated diffusion tensor map was smoothed using a median filter nonlinear registration with the standard space MNI152, 1 mm^3^). For every subject, the average values of the DTI measures (i.e., FA, MA, and RD) were delimited using the regions LIST_REGIONS from the JHU-DTI atlas (Wakana et al., 2007).

### Statistical analysis of MRI data

The average volumetric (grey matter volume), surface-based (cortical complexity, sulci depth, gyrification index, cortical thickness), diffusion-based (fractional anisotropy, mode of anisotropy, radial diffusivity), and RS-fMRI (seeds: bilateral vmPFC, bilateral anterior ventral striatum (aVS), right insula, left dlPFC) for each of the atlas regions, and each participant’s total intracranial volume (TIV) values were used as input to a T-PLS regression (Lee et al., 2022) to predict individual differences in risk (log-transformed α) and intertemporal (log-transformed k) preferences. Demographic variables (age, sex, and IQ) were regressed out from the risk and intertemporal preference variables after log10-transformation.

We used an iterative cross-validation procedure that randomly sampled 10% (rounding down) of participants to test on, used the remaining 90% of participants to train the T-PLS model on, and repeated that procedure 1000 times for stable median out-of-sample predictions. We used random sampling instead of dividing participants into a set number of folds to make sure the prediction accuracy is not influenced by how participants are arbitrarily divided into folds. Within each cross-validation fold, the predictor was therefore trained on 95 and tested on 10 participants. Within each fold’s training data, we used a nested cross-validation procedure with the same random sampling approach (but only repeated 100 times to reduce computational cost) to select optimal tuning parameters, i.e., the number of PLS components (1 - 25) and features (ranging from 1 - 2263 features in steps of 10%) with highest prediction accuracy in terms of Pearson’s correlation. After the nested cross-validation, the T-PLS model was fitted to all training data using the optimal tuning curves to predict individual risk or intertemporal preferences for the left-out test data in that cross-validation fold. The final out-of-sample prediction accuracy is computed by taking the median Pearson’s correlations across all 1000 cross-validation test-folds.

To statistically evaluate if the out-of-sample prediction accuracies were better than chance, we ran the original analyses again but with randomly shuffled individual values for risk or intertemporal preferences. This procedure was repeated 5000 times to create a null-distribution to which we compared the median correlation values to compute a p-value.

T-PLS model coefficients were generated by building a separate predictor model based on the same random sampling procedure used for out-of-sample prediction (without nested cross-validation) to select optimal tuning parameters and fitting the T-PLS model to all participants’ data. To visualize the coefficients, we mapped them on to brain renderings according to the brain atlases.

## Results

We use the model-based preference measures log10(α) and log10(k) in our analyses below but have previously shown that these measures are strongly correlated with other model-based and model-free measures of preference in this dataset (Bergström et al., 2024; Jung et al., 2018). The results below are therefore likely to be robust with other measures of risk and intertemporal preferences. Moreover, both risk and intertemporal preferences had moderately high test-retest reliability in this sample. A subset of our sample (n = 85) was tested again 10 weeks later in a post-intervention session, and the correlation between the choice preferences in the baseline session we analyze below and 10 weeks later was r = 0.68 for risk and r = 0.71 for intertemporal preferences.

When using T-PLS with all multimodal brain measures together, we were able to predict individual differences in risk (log10α; Mdn r = 0.32, p = 0.009) and intertemporal (log10k, Mdn r = 0.31, p = 0.008) preferences (Figure 1A). Additionally, we compared the prediction achieved using the combination of all twelve structural properties to that achieved using each of the twelve brain structure properties individually. We found that combining all twelve brain properties produced higher prediction accuracy than the strongest individual brain measure for intertemporal preferences and the second highest prediction accuracy (after sulci depth) for risk preferences. The predictor models with combined measures thus had (on average) the highest prediction accuracy, and were the only ones better than chance, across both choice preferences (Figure 1D-E).

**Figure 1.**
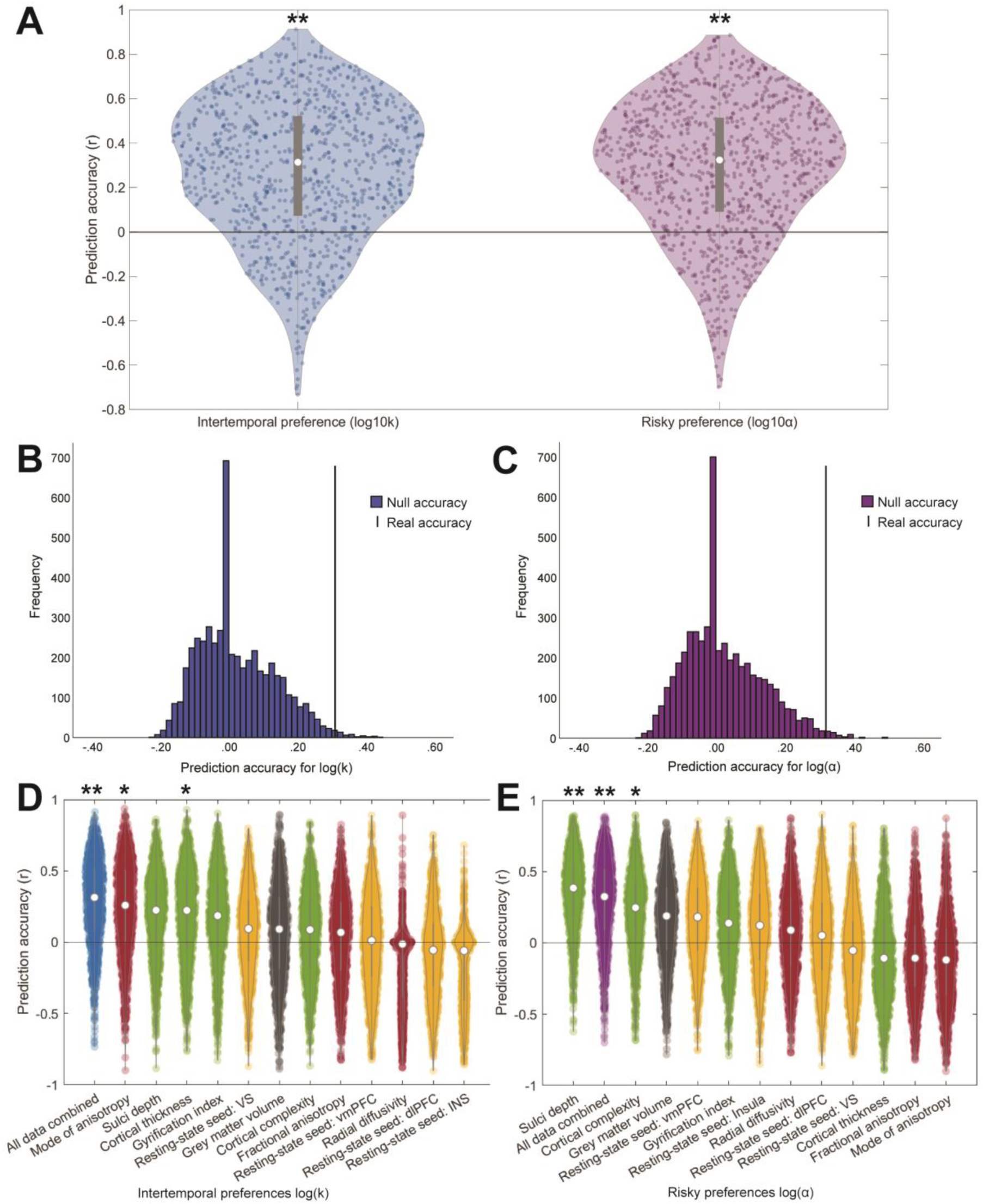
Out-of-sample prediction accuracy for risky and intertemporal preferences. (**A**) Violin plots showing out-of-sample prediction rate for intertemporal preferences (log10k) and risky preferences (log10α), using all twelve whole-brain structural properties. (**B**) Null distribution for intertemporal preferences (black line is real prediction accuracy. (**C**) Null distribution for risky preferences (black line is real prediction accuracy). (**D**) Violin plots showing prediction accuracy for risky preferences when using all combined properties and each property separately. (**E**) Violin plots showing prediction accuracy for risky preferences when using all combined properties and each property separately. Violin plots: The white dots show median out-of-sample prediction rate (Pearson’s r) for 1000 randomly sampled test data consisting of 10 participants each, the violin plot’s shape and colored dots show the distribution and prediction rate of each random sample, and the grey bars indicate the interquartile range. The black line indicates r = 0, * = p < 0.05, ** = p ≤ 0.01.

Specifically, it was possible to predict intertemporal preferences when only using mode of anisotropy (Mdn r = 0.26, p = 0.024) or cortical thickness (Mdn r = 0.22, p = 0.049), but not when using sulci depth (Mdn r = 0.22, p = 0.051), gyrification index (Mdn r = 0.19, p = 0.082), RS-fMRI seed: VS (Mdn r = 0.09, p = 0.220), grey matter volume (Mdn r = 0.09, p = 0.242), cortical complexity (Mdn r = 0.09, p = 0.248), fractional anisotropy (Mdn r = 0.07, p = 0.296), RS-fMRI seed: vmPFC (Mdn r = 0.01, p = 0.445), radial diffusivity (Mdn r = -0.01, p = 0.533), RS-fMRI seed: mid-dlPFC (Mdn r = -0.06, p = 0.673), or RS-fMRI seed: insula (Mdn r = -0.06, p = 0.677).

It was possible to predict risk preferences when only using sulci depth (Mdn r = 0.38, p = 0.001) or cortical complexity (Mdn r = 0.25, p = 0.03), but not when using GMV (Mdn r = 0.19, p = 0.083), RS-fMRI seed: vmPFC (Mdn r = 0.18, p = 0.089), gyrification index (Mdn r = 0.14, p = 0.142), RS-fMRI seed: insula (Mdn r = 0.12, p = 0.178), radial diffusivity (Mdn r = 0.09, p = 0.244), RS-fMRI seed: mid-dlPFC (Mdn r = 0.05, p = 0.351), RS-fMRI seed: VS (Mdn r = -0.05, p = 0.660), fractional anisotropy (Mdn r = -0.11, p = 0.859), cortical thickness (Mdn r = -0.11, p = 0.859), or mode of anisotropy (Mdn r = -0.12, p = 0.864).

To see the relative contribution of the different brain areas and brain properties to predicting risk and intertemporal preferences, we visualized the coefficients of the prediction model (Figure 2-5). Warm colors indicate a positive coefficient on the brain measure when predicting either a greater preference for risky rewards or a greater preference for immediate rewards, while cold colors indicate a negative coefficient when predicting the same behaviors. For both risky and intertemporal preferences, the T-PLS model used regions from all twelve structural brain properties. All brain properties thus contributed to the overall prediction. Based on the size of the coefficients, the largest relative contribution to both choice preferences came from GMV (Figure 2) and the surface-based measures sulci depth and gyrification index, while cortical complexity and cortical thickness had a more moderate contribution (Figure 3). RS-fMRI connectivity had a moderate contribution to risky preferences but a lesser contribution to intertemporal preferences (Figure 4). The smallest contribution came from diffusion-based properties (mode of anisotropy, fractional anisotropy, and radial diffusivity; Figure 5).

**Figure 2.**
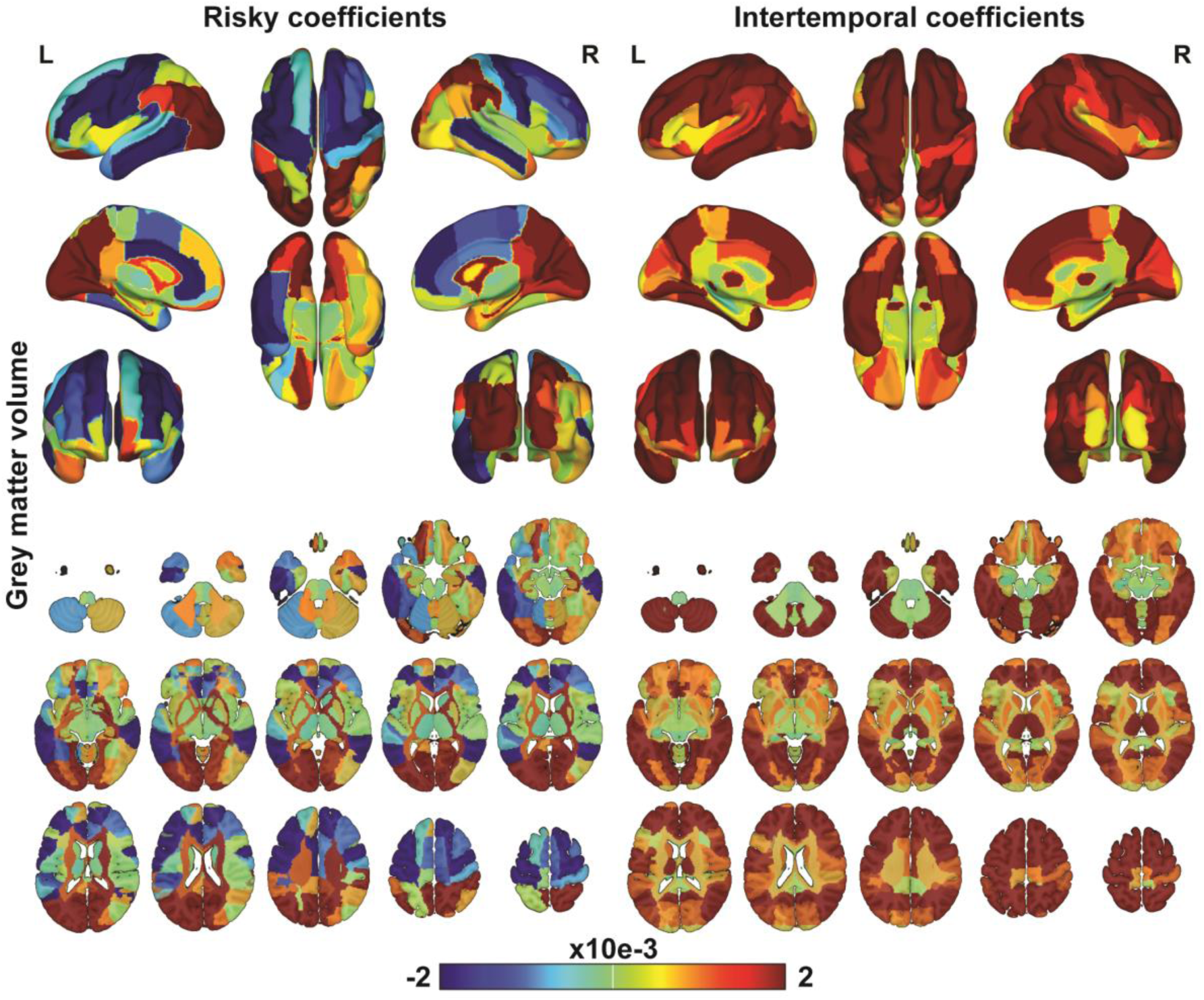
Grey matter volume coefficients. Thresholded Partial Least Square coefficients showing the relative contribution from grey matter volume in Neuromorphometric Atlas areas to the prediction rate for risky and intertemporal preferences.

**Figure 3.**
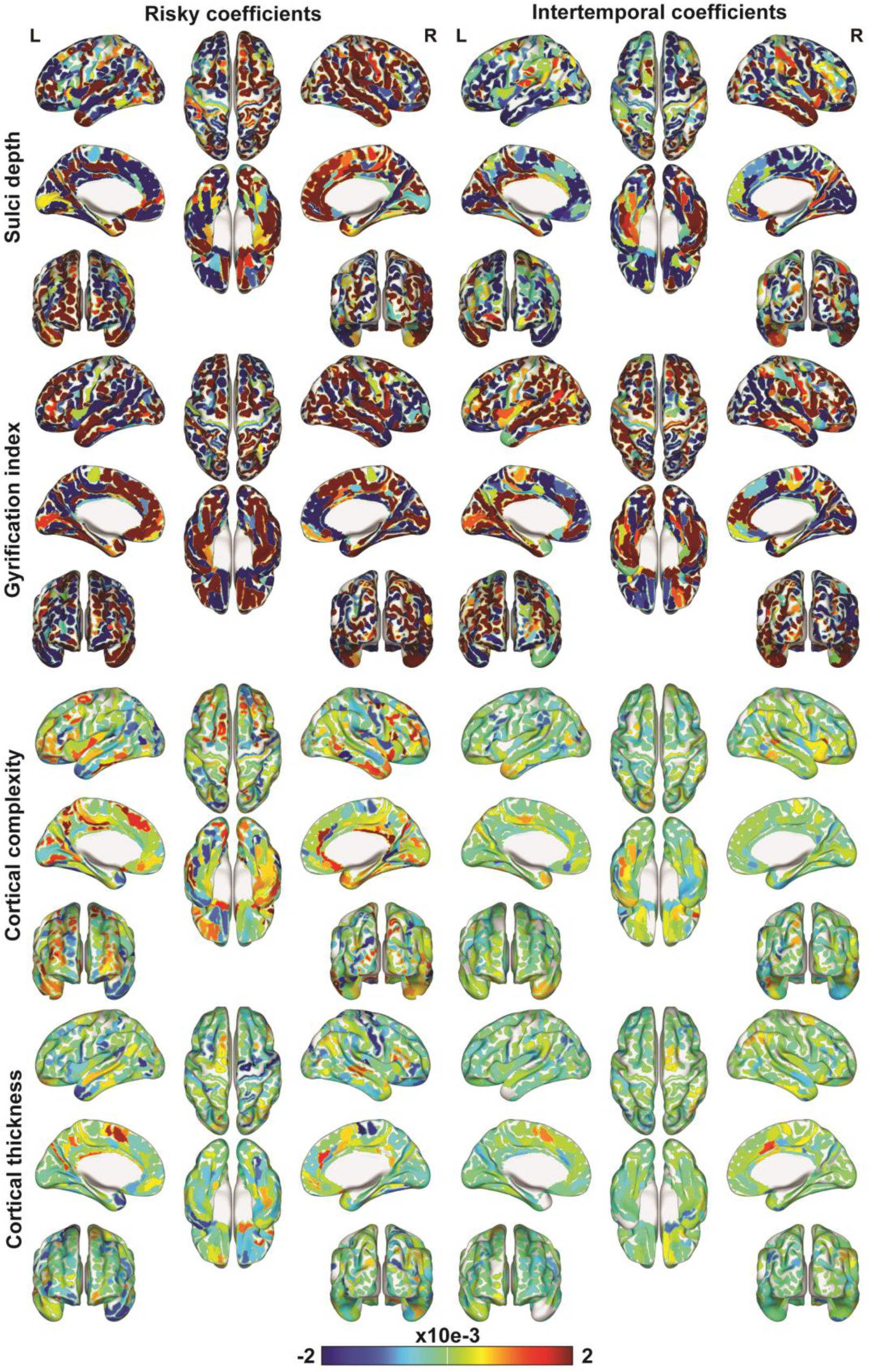
Surface-based coefficients. Thresholded Partial Least Square coefficients showing the relative contribution from surface-based measures in HCP-MMP surface atlas areas to the prediction rate for risky and intertemporal preferences.

**Figure 4.**
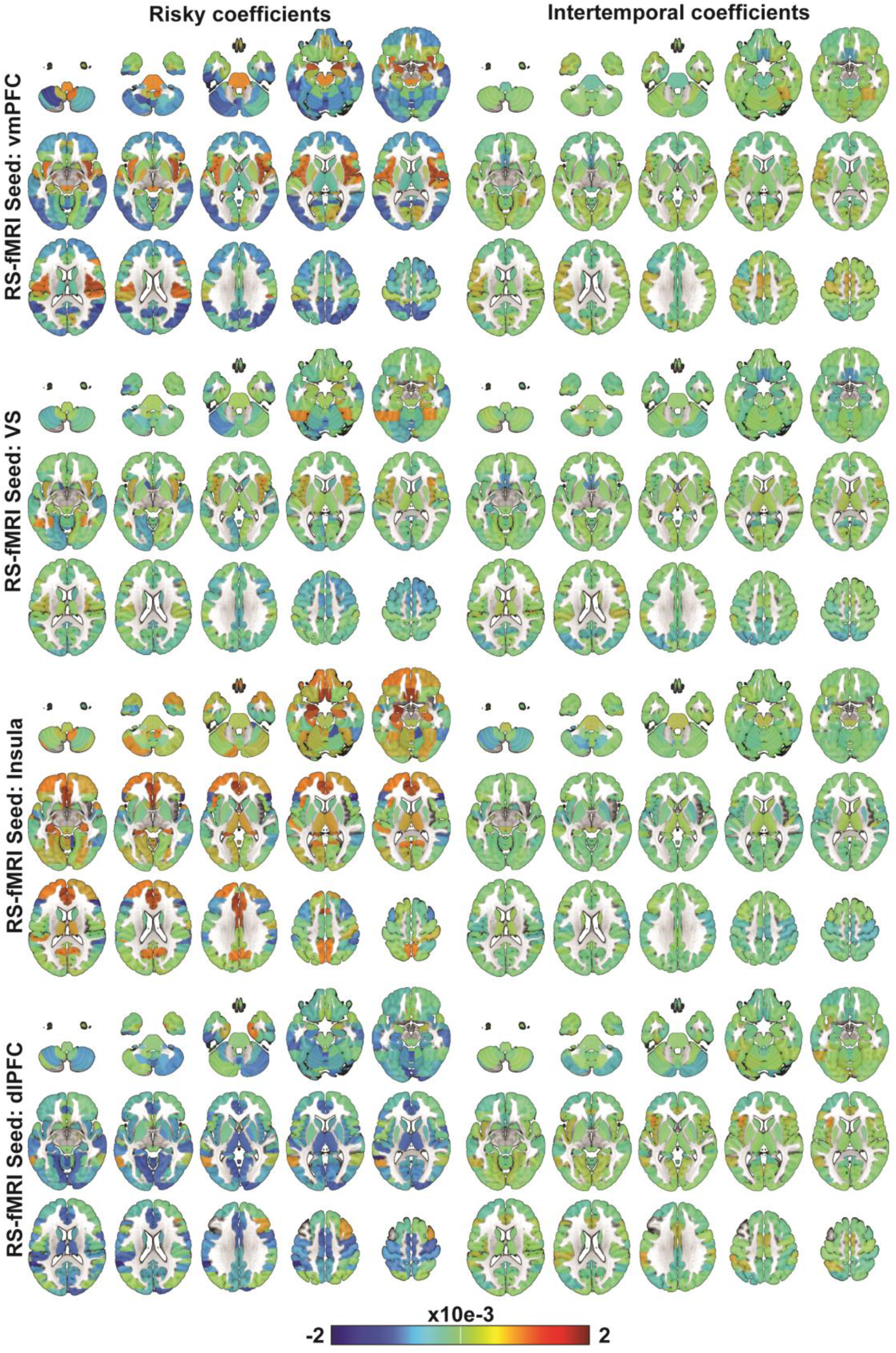
Resting-state fMRI coefficients. Thresholded Partial Least Square coefficients showing the relative contribution from resting-state fMRI in Conn toolbox atlas areas to the prediction rate for risky and intertemporal preferences.

**Figure 5.**
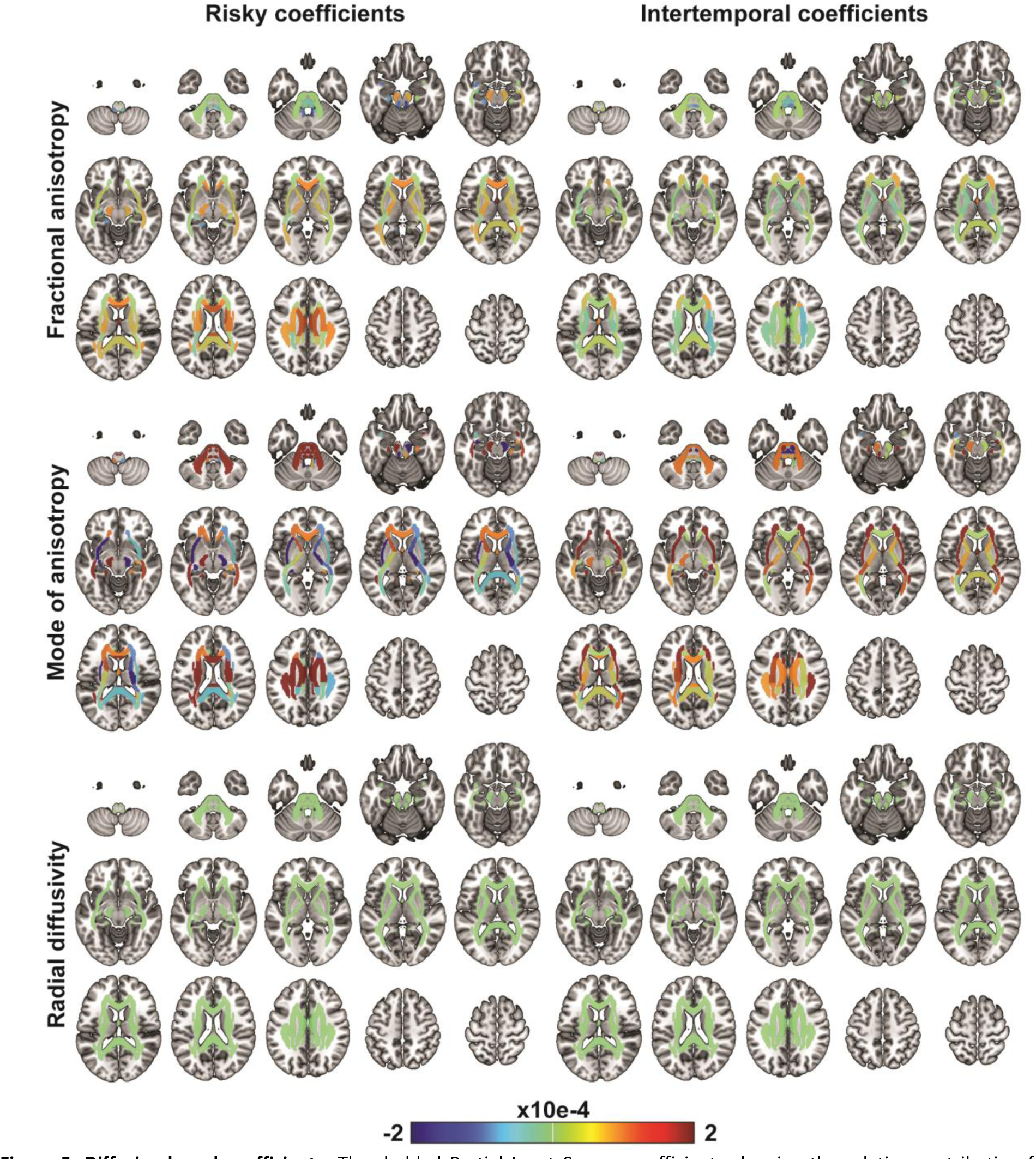
Diffusion-based coefficients. Thresholded Partial Least Square coefficients showing the relative contribution from diffusion-based measures in JHU Hammer Atlas areas to the prediction rate for risky and intertemporal preferences. Note that this figure has a different scale than the other figures to be able to visualize differences between brain regions. Note that the scale used for the diffusion-based coefficients is smaller than the other measures’ coefficients because otherwise it would be impossible to discern any internal difference between different diffusion measures or brain areas within the same diffusion measure.

## Discussion

We demonstrate a novel multivariate approach to predicting behavioral traits using a multimodal MRI data set. Our approach used the multivariate technique Thresholded Partial Least Squares (T-PLS; Lee et al., 2022) on twelve brain property measures (anatomical, RS-fMRI, and diffusion) from a multimodal MRI data set to predict individual differences in risk and intertemporal preferences. Predictions using all brain properties together were generally more accurate than those using any single brain property separately. Examining the coefficients in the combined predictor model, the largest contributions came from gyrification, sulci depth, and grey matter volume; with intermediate contributions from cortical complexity, cortical thickness, and resting-state connectivity; and the smallest contributions from diffusion-based measures. It is worth noting that the best prediction accuracy for the combined predictor model was achieved by using features across all brain measures despite the varying performance when using them separately, suggesting that all brain measures contained information that was useful for predicting the preferences.

The multivariate approach demonstrated here presents several advantages over the standard approach of performing univariate analyses on a single brain structural measure. First, this approach takes advantage of information from the whole brain instead of only a single voxel or atlas region. Second, this approach also considers information across multiple brain measures instead of only one measure. Third, this approach tests if the predictions generalize to new individuals instead of only demonstrating a relationship within a group – which does not necessarily translate to significant out-of-sample predictions. At the same time, the interpretability of T-PLS coefficients can inform us about the relative contribution of different brain properties from different brain regions to prediction. Our results here are not only an existence of proof for the potential power of this approach for examining relationships between brain structure and individual trait differences, but they also have wider implications across basic, translational, and clinical research fields. For example, this methodological approach may be useful to improve clinical diagnosis predictions and give a better understanding of the intricate relationships between brain properties and brain areas that underlie different clinical disorders. Though there are cases where multimodal brain structure data has been used for predictive diagnostics, it seems to be a minority of cases and often relatively few brain measures and small sample sizes (for review, see Mateos-Pérez et al., 2018). With larger sample size, combining more brain structure measures becomes more feasible and may improve diagnostic accuracy and insight into underlying brain structure.

It is difficult to benchmark our out-of-sample prediction accuracies for risky or intertemporal preferences against other studies. In a recent study, Koban et al. (2023) used task-based fMRI markers to predict individual differences in intertemporal preferences with a higher prediction accuracy than ours. However, studies in other domains have found similar prediction accuracies as ours when, for example, using RS-fMRI connectivity to predict individual differences in creative ability (Beaty et al., 2018) or cross-decoding individual differences in sustained attention from task-based fMRI to RS-fMRI connectivity (Rosenberg et al., 2016). Within-sample correlations of functional or structural brain measures of single brain areas have had similar or weaker correlations than ours (e.g., Cooper et al., 2013; Jung et al., 2018; Quan et al., 2022), but within-sample correlations do not necessarily generalize to equally high or even significant out-of-sample prediction accuracy. It is possible that task-based fMRI data may provide more accurate predictions than structural brain measures, especially when collected during the behavior of interest; function may be more closely linked to individual differences than structure. However, the strength and interest in prediction from brain structure measures lies in their practical application, including that because these measures are not specific to any given task, the same data set can be used to predict any number of cognitive functions or behavioral traits (Kable & Levy, 2015).

When used individually for prediction, none of the single brain measures significantly predicted both risk and intertemporal preferences. Sulci depth was closest, with the highest prediction accuracy for risk preferences and almost reaching statistical significance for predicting intertemporal preferences. This may be because these brain measures have a relatively low signal-to-noise ratio and/or because the most relevant brain measure differs across behaviors. The multivariate multimodal approach used here can increase prediction accuracy by including many potentially relevant measures and combining measures to increase signal-to-noise.

Interestingly, we observed a rough hierarchy in terms of the relative contribution of the different brain structural measures to predicting individual preferences. Gyrification, sulci depth, and GMV made the strongest contributions to predicting both risk and intertemporal preferences, with cortical complexity and cortical thickness making more moderate contributions. RS-fMRI connectivity made a moderate contribution that was stronger when predicting risk than intertemporal preferences. Diffusion measures made the smallest relative contribution to predicting both preferences. An interesting question for future research will be whether this rough hierarchy holds across other kinds of structural, connectivity and diffusion measures, as well as across other kinds of individual differences.

The novel approach of using a multivariate technique on multimodal brain structure properties makes it more challenging to relate our results to previous univariate work given the added complexity. However, consistently with previous literature, we found a positive relationship between a preference for larger risky rewards and GMV in the right PPC (Gilaie-Dotan et al., 2014b; Grubb et al., 2016b; Quan et al., 2022), GMV in (right) amygdala and amygdala-vmPFC RS-fMRI connectivity in the same data set (Jung et al., 2018); and a positive relationship between a preference for smaller immediate rewards and GMV in vmPFC, ACC, VS, and PCC (Cho et al., 2013; Schwartz et al., 2010), and a negative relationship between a preference for smaller immediate rewards and cortical thickness in vmPFC (Bergström et al., 2024; Bernhardt et al., 2014; Drobetz et al., 2014; Pehlivanova et al., 2018). We previously showed a negative relationship between a preference for larger risky and smaller immediate rewards and cortical complexity in vmPFC with partly the same data set (Bergström et al., 2024), but we did not see a similar result here. This discrepancy could be because the area of the voxel-wise cluster previously found was here divided into, and averaged within, several anatomically defined sub regions of vmPFC.

There is ongoing debate about the sample sizes needed to examine relationships between brain structure and behavior. Marek et al. argued that the typical (median) sample size for brain-wide association studies (n = 23) was orders of magnitude too small, and that sample sizes in the thousands are necessary to reliably link brain structure to behavior (Marek et al., 2022). However, the latter claim is not without controversy, as some argue that Marek et al.’s (2022) use of average estimates of power, replication rate, and statistical error across brain regions underestimate reproducibility of brain regions with stronger than average associations (Cecchetti & Handjaras, 2022), that they generalized their conclusions from unreasonably low effect sizes and that a more reasonable sample size may be around 200 participants for univariate analysis (DeYoung et al., 2022). Importantly, it has been demonstrated that reproducible multivariate out-of-sample predictions can be achieved with around 100 participants when using structural brain data (Makowski et al., 2023). Given our sample size (n = 105), our results would support this conclusion.

In the current study, we averaged our brain measures within brain atlas regions to reduce what otherwise would be a few million features to only 2263 features for reduced dimensionality and computational feasibility. Future work could examine the impact of this decision. It is possible that we lost signal by reducing the spatial resolution of the brain measure by averaging features within relatively large brain atlas regions. Alternatively, it is possible that such averaging increased the signal to noise ratio of the measures. It is also possible that larger data sets to train the models on may further increase prediction accuracy. For the current study, we chose an approach that achieved adequate spatial resolution while allowing the inclusion of multiple different brain measures covering the whole brain, given our sample size.

In conclusion, we demonstrate that a novel multivariate approach using multimodal MRI data can successfully predict individual differences in risk and intertemporal preferences, and that combining different brain measures using this approach can improve prediction accuracies, while still permitting an examination of the intricate relationships between different brain properties, brain areas, and behavioral preferences. This approach may also be well suited for use basic, translational, and clinical research fields.

## Data and code availability

MRI and behavioral data are available at OpenNeuro (https://doi.org/10.18112/openneuro.ds002843.v1.0.1). MATLAB code for estimating risk and intertemporal preferences is available at GitHub (https://github.com/sangillee/UMm).

## Author Contributions

FB: Conceptualization (lead), funding acquisition, methodology, validation, formal analysis, investigation, data curation, writing – original draft, writing – review & editing, and visualization. GG: Methodology and writing – review & editing. SL: Methodology and writing – review & editing. CL: Conceptualization (supporting), funding acquisition, and writing – review & editing. JWK: Conceptualization (supporting), funding acquisition, data curation, and writing – review & editing.

## Acknowledgements

This work was supported by Fundação para a Ciência e Tecnologia (CEECIND/03661/2017 to F.B. and Advanced Computing Project 2021.09640.CPCA platform Navigator to F.B. and J.W.K.); and National Cancer Institute Grants (R01-CA-170297 to J.W.K. and C.L. and R35-CA-197461 to C.L.). The authors acknowledge Fernanda Martinez for programming assistance, and the Laboratory for Advanced Computing at University of Coimbra for providing high performance computing resources that contributed to the research results reported within this paper. URL: https://www.uc.pt/lca.

